## Supplementary Information for "Cyclodipeptide oxidase is an enzyme filament"

### Table of Contents

| | |
|---|---|
| Supplementary Fig. 1. Sequence similarity network analysis of CDO-encoding gene clusters | 3 |
| Supplementary Fig. 2. Stability screens of AlbAB filaments | 5 |
| Supplementary Fig. 3. Saturation kinetics and assembly analysis of AlbAB and AlbB | 6 |
| Supplementary Fig. 4. Cryo-EM data collection and processing workflow | 7 |
| Supplementary Table 1. Cryo-EM data collection, refinement, and validation statistics | 8 |
| Supplementary Fig. 5. Local resolution estimation of the AlbAB cryo-EM map | 9 |
| Supplementary Fig. 6. Map-to-model fit of the AlbAB cryo-EM density | 10 |
| Supplementary Fig. 7. Structural analysis of AlbA and comparison to other NTR-like proteins | 11 |
| Supplementary Fig. 8. AlbB structure analysis | 13 |
| Supplementary Fig. 9. Sequence conservation in CDOs | 14 |
| Supplementary Fig. 10. Interactions and sequence conservation at the active site | 16 |
| Supplementary Fig. 11. Investigating the interaction of AlbAB with AlbC and tRNA | 18 |
| Supplementary Table 2. Protein sequences of constructs used in this study | 19 |
| References | 20 |

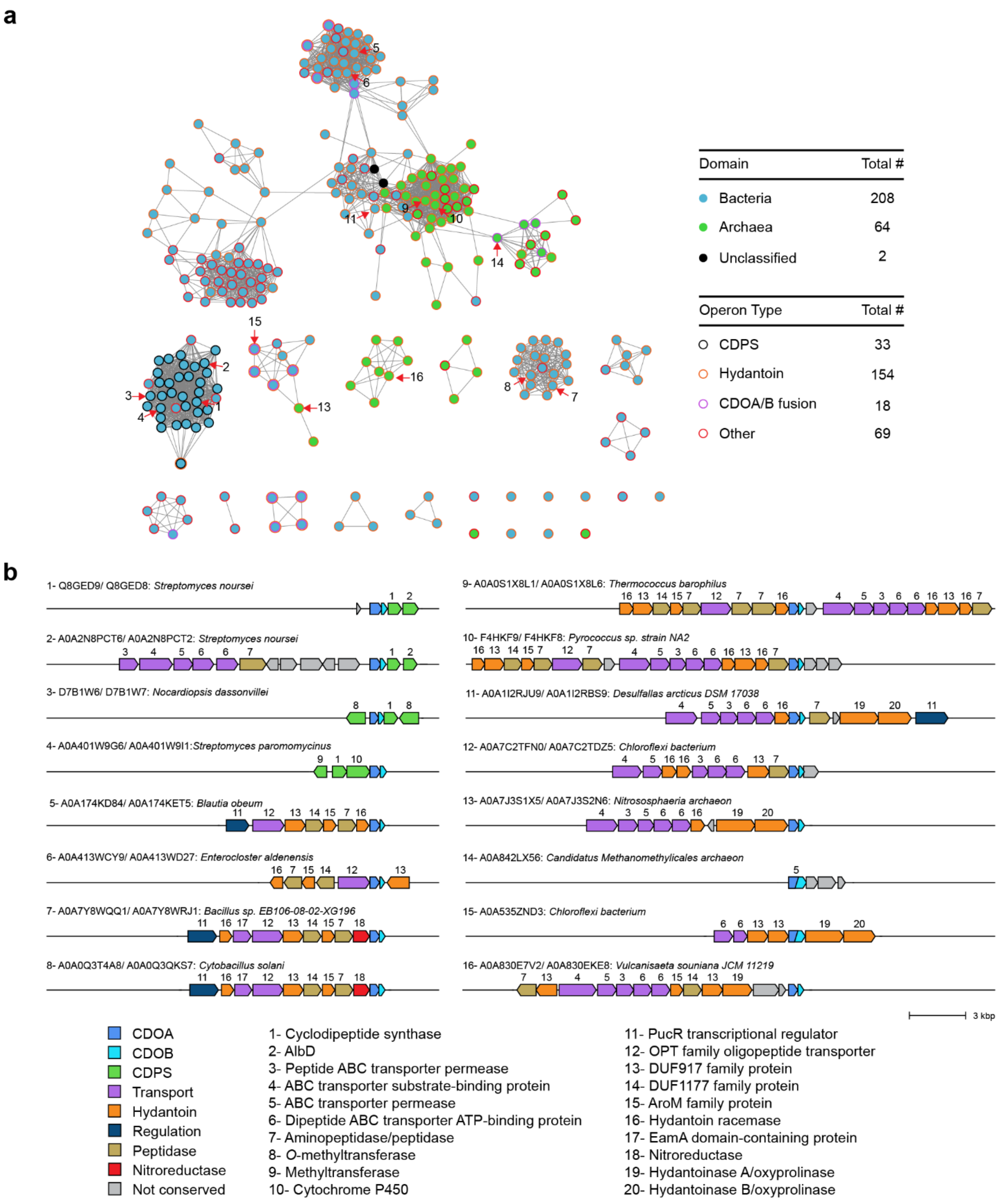

**Supplementary Fig. 1: Sequence similarity network analysis and diversity of CDO-encoding gene clusters. a**, Sequence similarity network (SSN) analysis shows the similarity between clusters of concatenated CDOA and CDOB sequences with an identity threshold of 46%. 274 total operons were identified with complete amino acid sequences of both the CDOA and the CDOB from 64 archaeal (green nodes) and 208 bacterial genomes (blue nodes). Two CDOs from unclassified metagenomic samples (black nodes) were also identified. 33 Actinomycetota gene clusters containing CDPS-encoding genes (black outline) were contained in a single cluster. 154 gene clusters were identified to encode proteins associated with hydantoin/allantoin modification (orange outline). 18 gene clusters were found to encode CDOA/B fusion proteins (purple outline). The SSN was generated using EFI-EST[1] and annotated using CytoScape v3.1.0[2]. **b**, The operon structure and organization of CDO-encoding gene clusters is diverse. Representative gene clusters 1-16 correspond to the nodes numbered in **a**. Conserved sets of genes are color coded according to predicted function and annotated below.

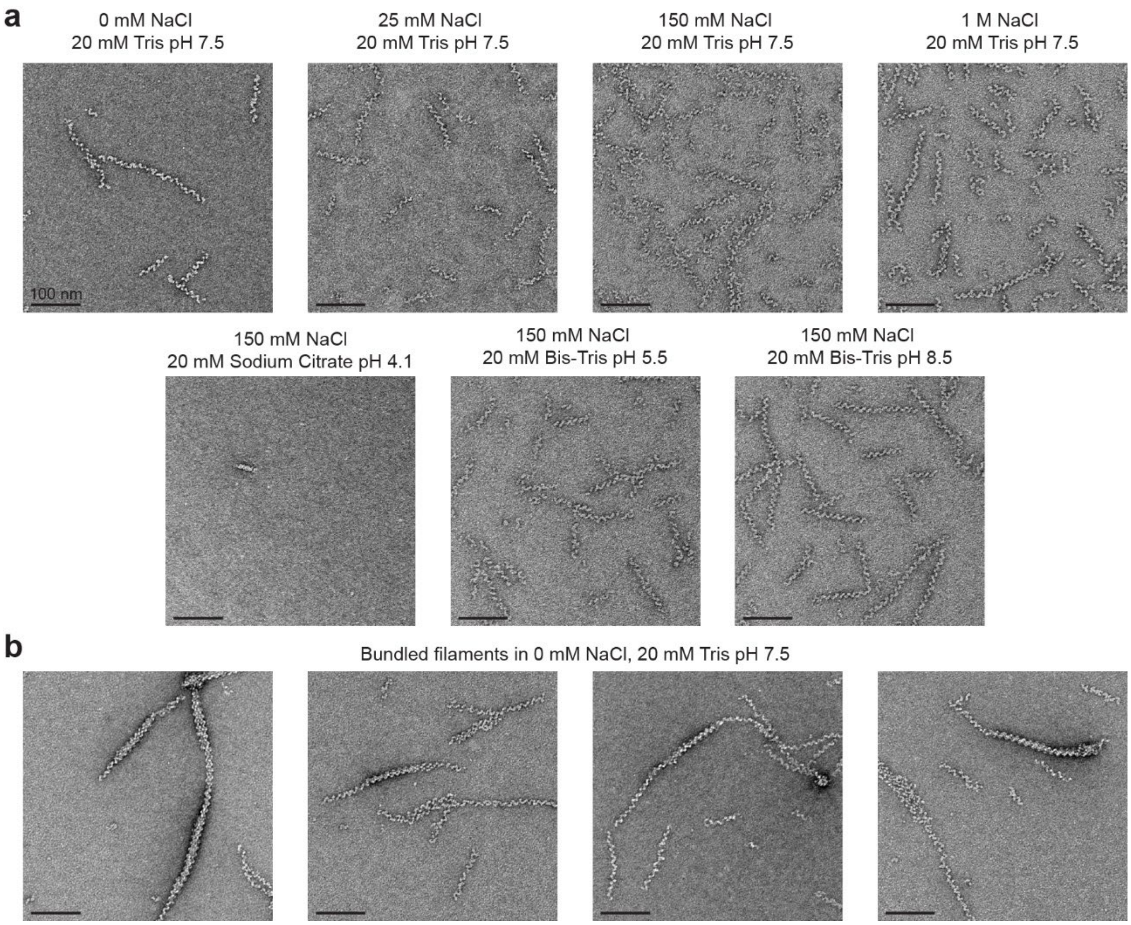

**Supplementary Fig. 2: Stability screens of AlbAB filaments. a**, AlbAB filaments were dialyzed against each of the conditions listed at 4°C for 18 h and subsequently analyzed by negative stain TEM. The AlbAB filaments were stable at pH 7.5 from 0 to 1 M NaCl, and at pH 5.5 to 8.5 in 150 mM NaCl. Filaments were not observed when dialyzed against 150 mM NaCl, 20 mM sodium citrate pH 4.1, suggesting they may aggregate or depolymerize below pH 5.5. **b**, A small fraction of purified filaments were found to form bundles of two or more filaments. Bundles were observed in conditions up to 150 mM NaCl. All scale bars correspond to 100 nm.

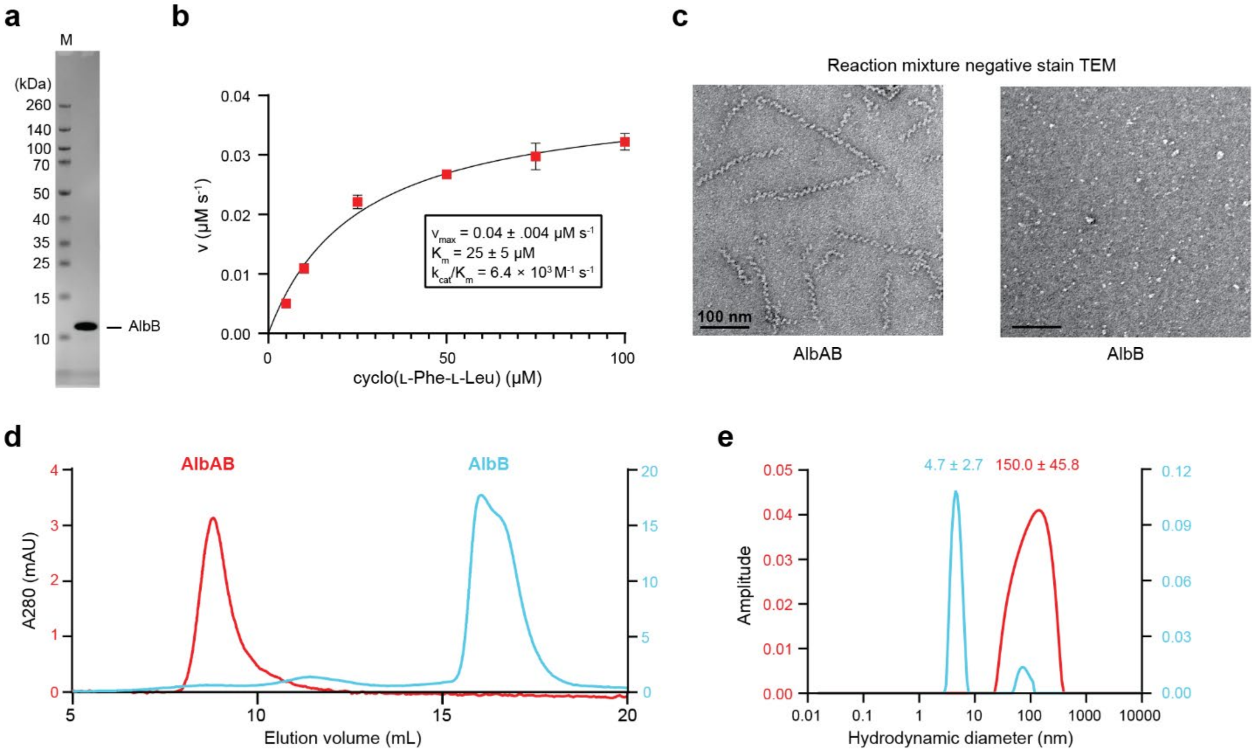

**Supplementary Fig. 3: Saturation kinetics and assembly analysis of AlbAB and AlbB. a**, SDS-PAGE analysis of purified AlbB. **b**, Saturation kinetics fit with a standard Michaelis-Menton curve of filamentous AlbAB under steady-state conditions. **c,** Negative stain TEM micrographs of AlbAB and AlbB in assay conditions consisting of 0.5 μM protein, 100 mM Tris pH 7.5, 0.1 U horseradish peroxidase, 1 mM 4-HPA. 100 μM cyclo(L-Phe-L-Leu) was added 30 seconds prior to staining the sample on the grid. This confirms that the catalytically active form of AlbAB are enzyme filaments. **d**, Representative Superdex S-200 elution profiles of purified AlbAB and AlbB. AlbAB elutes in the void volume due to the high mass and elongated dimensions of the filaments. AlbB elution corresponds to a dimeric assembly (23.1 kDa). **e**, DLS analysis of AlbAB agrees with TEM analysis and shows filaments to have a broad distribution of hydrodynamic radii with a peak diameter of 150.0 ± 45.8 nm. The major peak of AlbB has a hydrodynamic radius with a diameter of 4.7 ± 2.67 nm and a predicted molecular weight of 23.0 kilodaltons, further supporting a dimeric assembly in solution.

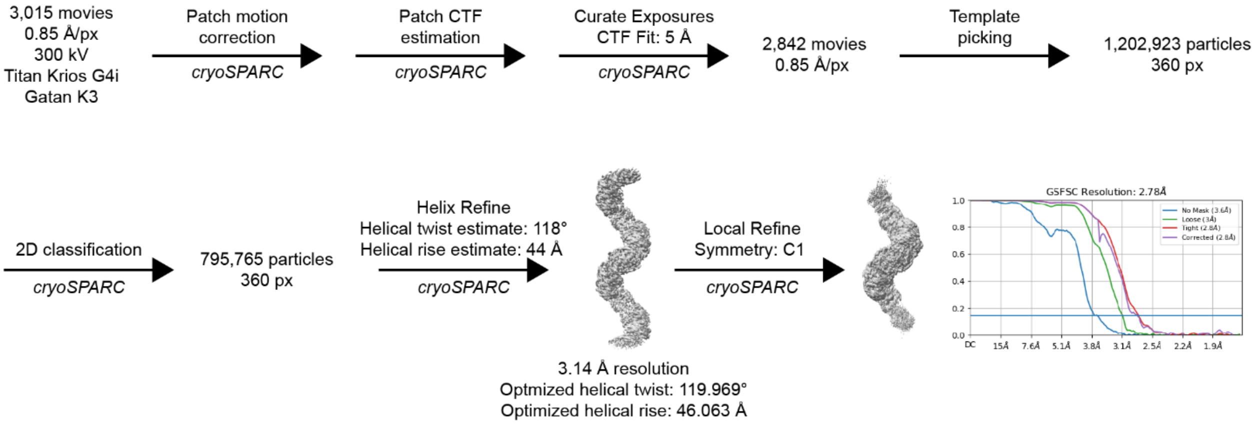

**Supplementary Fig 4: Cryo-EM data collection and processing workflow.** Data collection and processing workflow to generate the reconstructed volume for AlbAB. The final local refinement was used for atomic model building.

**Supplementary Table 1: Cryo-EM data collection, refinement, and validation statistics.**

| | AlbAB (EMDB-42114) (PDB ID: 8UC3) |
|---|---|
| **Data collection and processing** | |
| Magnification | 105,000x |
| Voltage (kV) | 300 |
| Electron exposure ($e^-/Å^2$) | 50.12 |
| Defocus range (μm) | -1.0 to -2.5 |
| Pixel size (Å) | 0.85 |
| Symmetry imposed | C1 (local refinement) |
| Initial particle images (no.) | 1,202,923 |
| Final particle images (no.) | 795,765 |
| Map resolution (Å) | 2.78 |
| FSC threshold | 0.143 |
| **Refinement** | |
| Initial model used | AlphaFold |
| Model resolution (Å) | 3.2 |
| FSC threshold | 0.5 |
| Map sharpening *B* factor ($Å^2$) | -107.4 |
| Model composition | |
| Non-hydrogen atoms | 4,304 |
| Protein residues | 556 |
| Ligands | 2 |
| *B* factors ($Å^2$) | |
| Protein | 66.66 |
| Ligands | 64.45 |
| r.m.s. deviations | |
| Bond lengths (Å) | 0.005 |
| Bond angles (°) | 1.111 |
| Validation | |
| MolProbity score | 2.06 |
| Clashscore | 12.15 |
| Poor rotamers (%) | 1.78 |
| Ramachandran plot | |
| Favored (%) | 95.99 |
| Allowed (%) | 4.01 |
| Disallowed (%) | 0 |

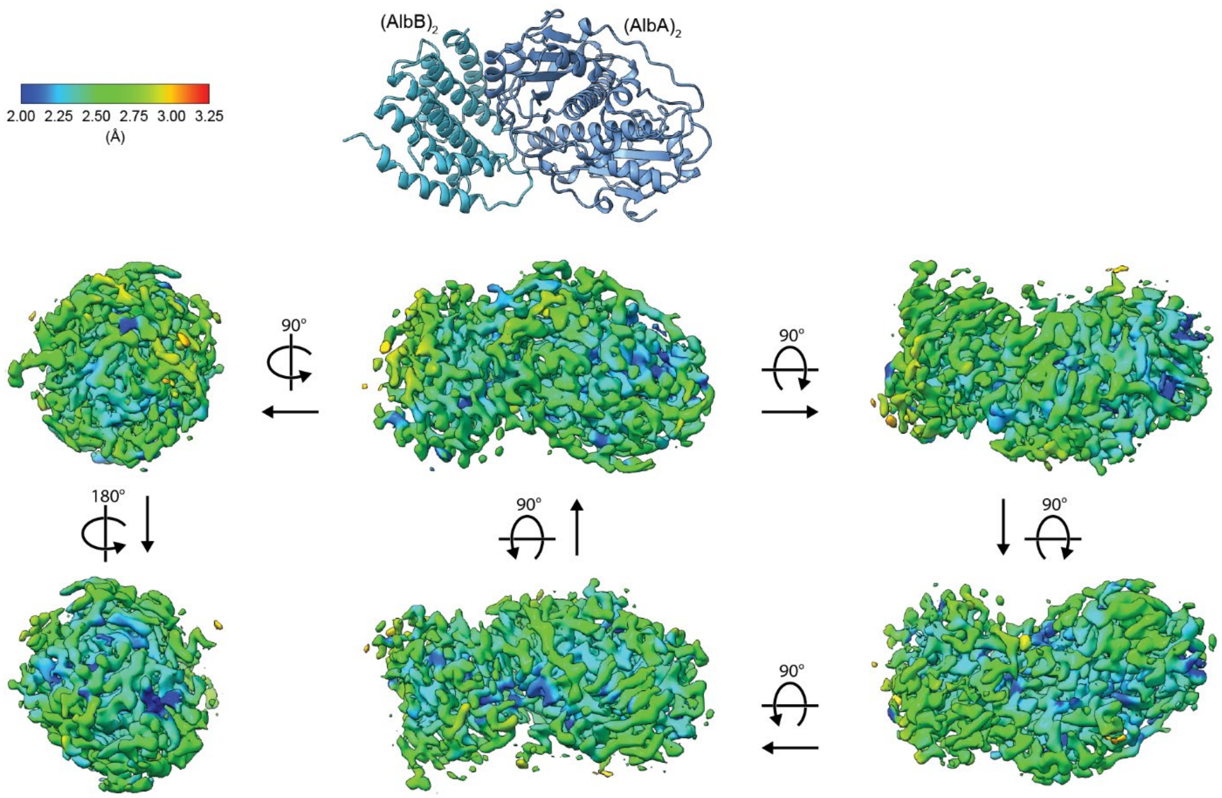

**Supplementary Fig. 5: Local resolution estimation of the AlbAB cryo-EM map.** Different perspectives of the C1 local refinement cryo-EM map surrounding a single AlbAB heterotetramer colored by estimated local resolution.

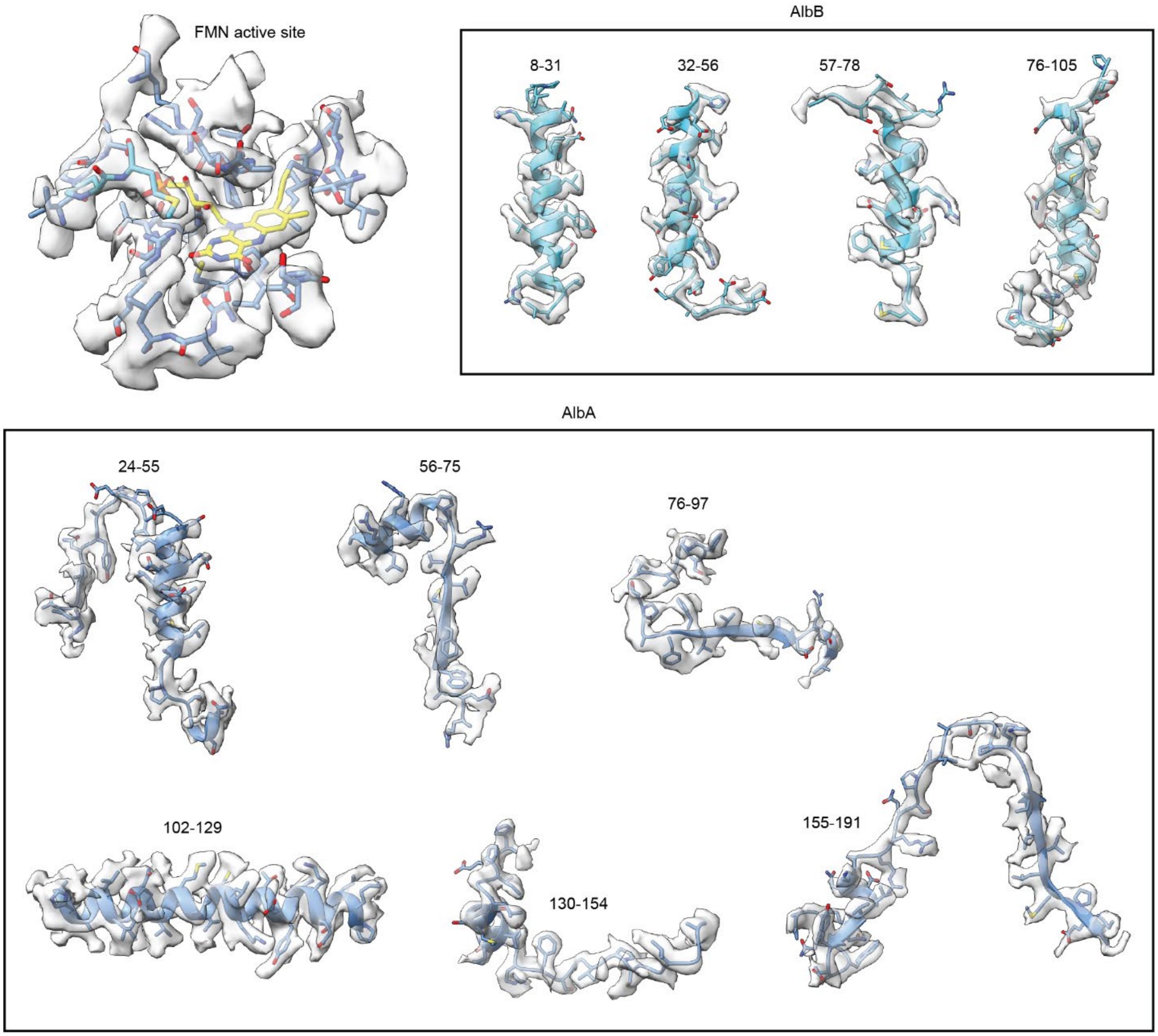

**Supplementary Fig. 6: Map-to-model fit of the AlbAB cryo-EM density.** Representative densities covering the majority of the sequences for AlbA and AlbB, including the area surrounding the FMN active site. Atomic models are shown as well to demonstrate map-to-model fit. The respective sequence numbering is shown.

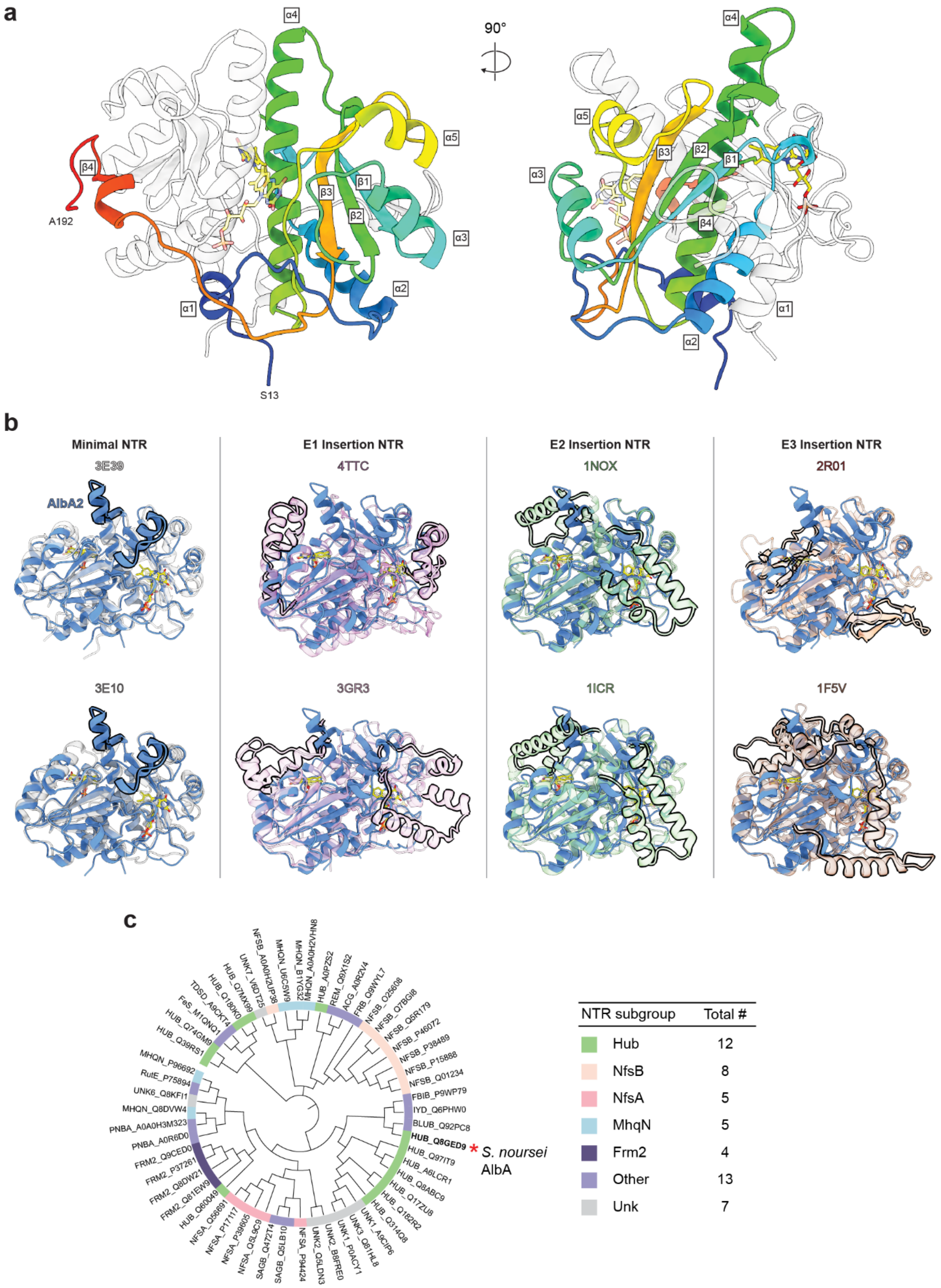
a
α4
90°
α5
β4
A192
β3
β1
β2
α3
α2
α1
S13
β2
β1
b
Minimal NTR
3E39
AlbA2
3E10
E1 Insertion NTR
4TTC
3GR3
E2 Insertion NTR
1NOX
1ICR
E3 Insertion NTR
2R01
1F5V
c
HUB_Q8GED9
* S. noursei AlbA
NTR subgroup
Total #
Hub 12
NfsB 8
NfsA 5
MhqN 5
Frm2 4
Other 13
Unk 7

**Supplementary Fig. 7: Structural analysis of AlbA and comparison to other NTR-like proteins. a**, AlbA is a dimer with a minimal nitroreductase (NTR)-like fold. An AlbA monomer in ribbon representation is shown colored as rainbow (N: blue to C: red). $\alpha$-helices and $\beta$-strands are labeled. **b**, Structural comparison of AlbA and other nitroreductases and nitroreductase-like proteins with determined structures. The proteins are grouped based on "hub" NTRs, NTRs containing an E1 insertion between alpha helices 1 and 2, NTRs containing an E2 insertion between beta strand 2 and helix 4, and NTRs containing an E3 extension located near the C-termini of representative proteins. When compared to the minimal NTR, AlbA has a slightly elongated E2 extension and does not contain an alpha helix where helix 4 should be in most proteins with an NTR-like fold. **c**, Phylogenetic tree comparing the amino acid sequence of AlbA with 54 other characterized NTRs representative of other NTR subgroups.

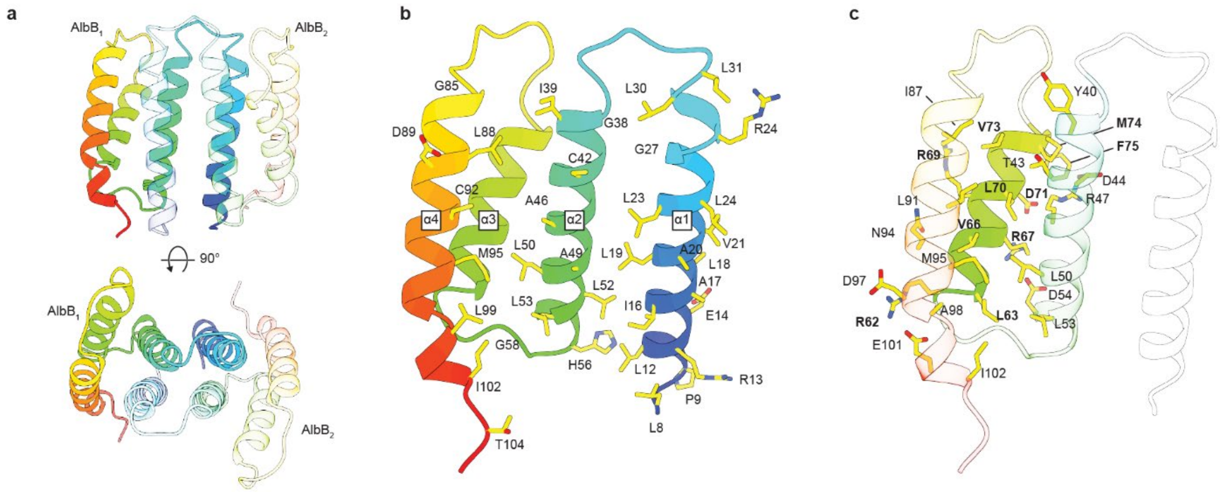

**Supplementary Fig. 8: AlbB structure analysis. a**, The $AlbB_2$ dimer as seen from two different perspectives. The chain of $AlbB_1$ is shown in rainbow coloring (N: blue to C: red). **b**, The AlbB monomer consists of four alpha-helices in an anti-parallel configuration. The 34 residues that contribute to the dimeric interface are labeled. **c**, Helix 3 does not contribute to the dimeric interface, but instead interacts intramolecularly with helices 2 and 4. Interacting residues at the interface between helix 3 and helices 2 and 4 are labeled, with helix 3 residues shown in bold.

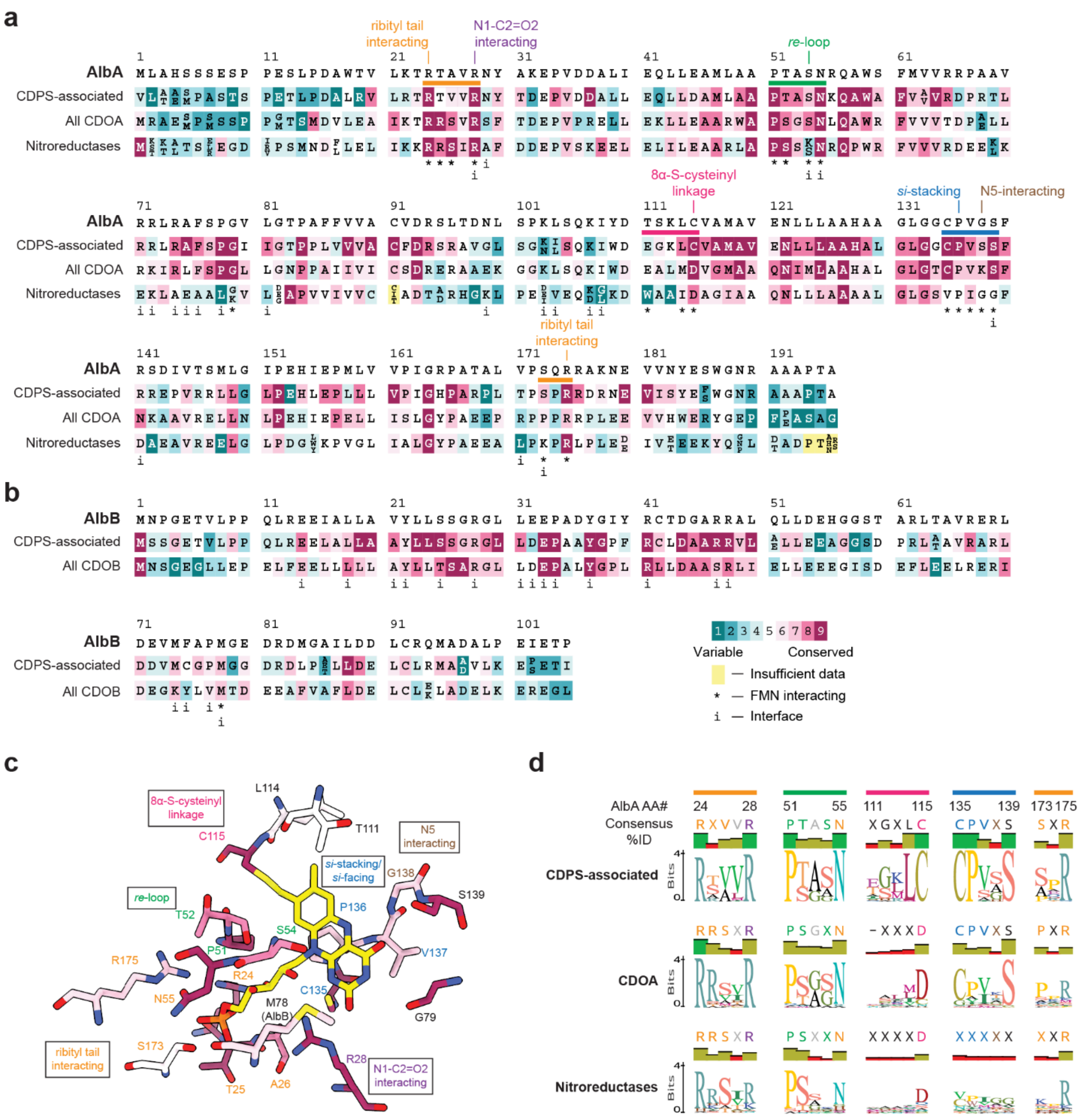

**Supplementary Fig. 9: Sequence conservation in CDOs. a**, Sequence conservation of AlbA according to ConSurf. The AlbA amino acid sequence is compared with all other CDPS-associated CDOA proteins, all CDOA proteins (including non-CDOPS-associated ones), and with 54 other NTRs used to assemble the phylogenetic tree in Supplementary Fig. 7. Amino acids are labeled by conservation score, contribution to FMN interactions, and contribution to the dimer-dimer interface in the AlbAB filament. **b**, Sequence conservation of AlbB according to ConSurf. The AlbB amino acid sequence is compared to all other CDPS-associated CDOB proteins and all other CDOB proteins (including non-CDPS-associated ones). Amino acids are labeled based on the same criteria as in **a**. **c**, FMN-interacting residues from AlbA and AlbB. Residues are color coded and labeled according to how they interact with FMN. **d**, Sequence logos representing the conservation at selected FMN-interacting residues of AlbA from panel **a**.

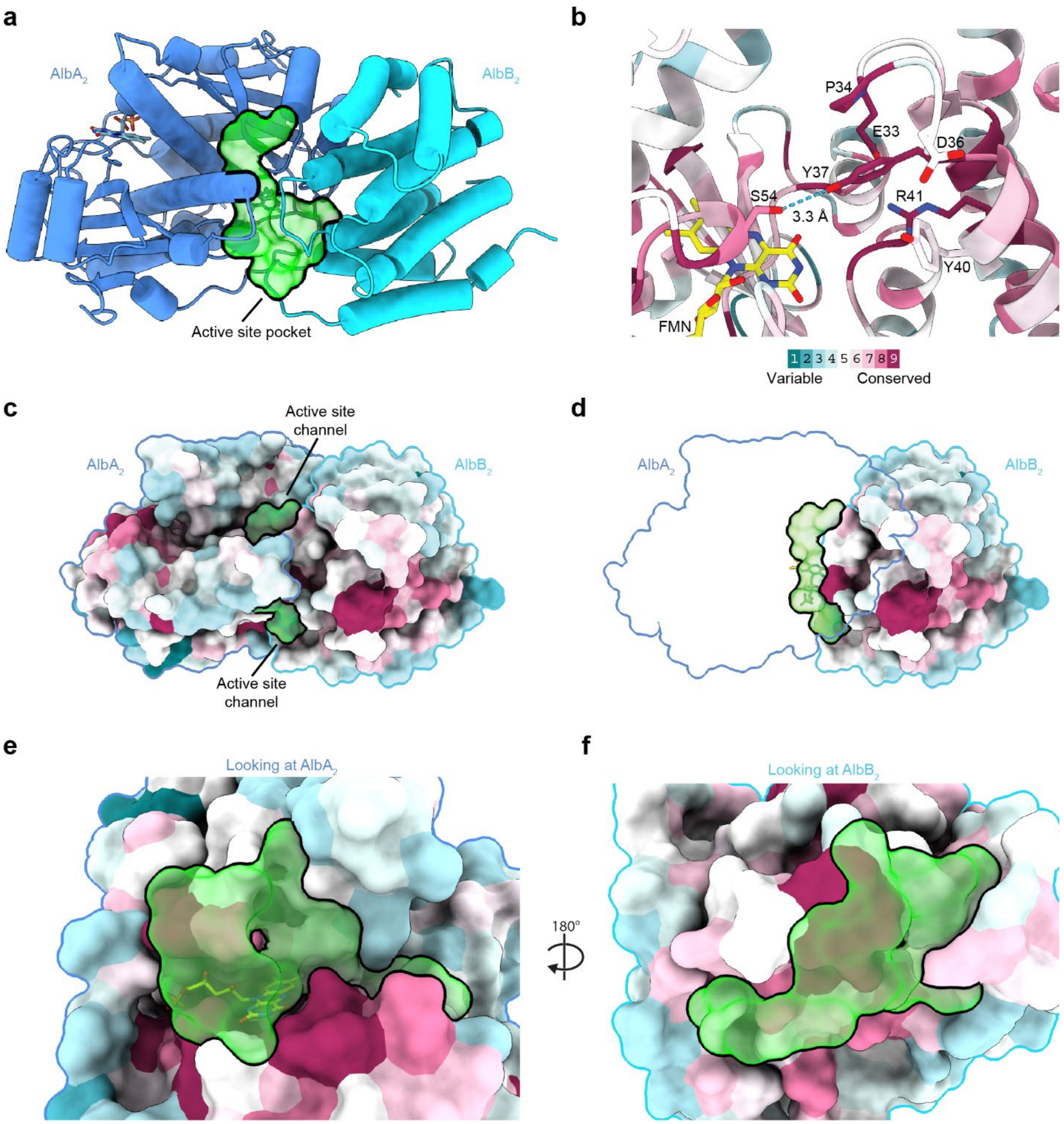

**Supplementary Fig. 10: Interactions and conservation at the active site.** **a**, An overview of the $AlbA_2$-$AlbB_2$ heterotetramer with the active site cavity – as calculated by FPocketWeb V1.0.1 (https://durrantlab.pitt.edu/fpocketweb/) – represented as a green surface. **b**, Conserved residues S54 and Y37 form a hydrogen bond above the *re* face of FMN. Other active site residues from AlbB within 5 Å of Y37 are also labeled. Residues are color coded by conservation according to ConSurf[3]. **c**, Same perspective as in **a**, surface representation colored by conservation highlighting the two access points to the active site. **d**, Same view as in **c** with AlbA completely transparent to reveal the connected active site cavity (green surface). **e**, View looking at the $AlbA_2$ dimer highlighting the active site cavity (green surface). **f**, View looking at the $AlbB_2$ dimer highlighting the active site cavity (green surface).

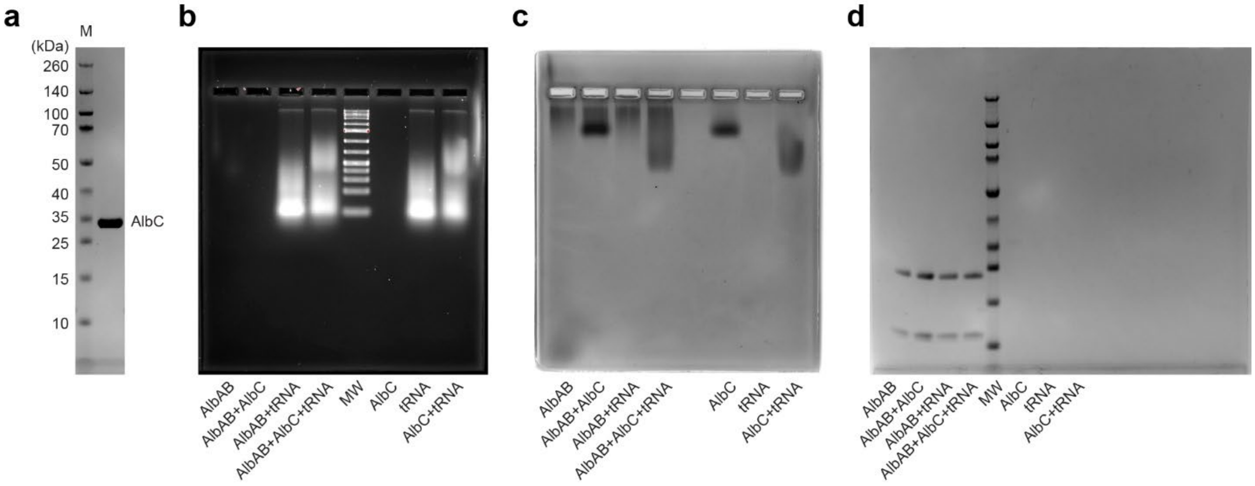

**Supplementary Fig. 11: Investigating the interaction of AlbAB with AlbC and tRNA.** **a**, SDS-PAGE gel of purified AlbC used in interaction studies. **b**, Full gel of the agarose gel shift assay shown in Fig. 6 stained with the nucleic acid stain GelCode Red. **c**, Full gel of the agarose gel shift assay shown in Fig. 6 stained with the protein stain GelCode Blue. **d**, Full SDS-PAGE gel of the void fraction analysis shown in Fig. 6.

**Supplementary Table 2: Protein sequences of constructs used in this study.**

| Name | Protein Sequence |
|---|---|
| AlbAB | MLAHSSSESPPESLPDAWTVLKTRTAVRNYAKEPVDDALIEQLLEAMLAAPTASNRQAWSFMVVRRPAAVRRLRAFSPGVLGTPAFFVVACVDRSLTDNLSPKLSQKIYDTSKLCVAMAVENLLLAAHAAGLGGCPVGSFRSDIVTSMLGIPEHIEPMLVVPIGRPATALVPSQRRAKNEVVNYESWGNRAAAPTA*MNPGETVLPPQLREEIALLAVYLLSSGRGLLEEPADYGIYRCTDGARRALQLLDEHGGSTARLTAVRERLDEVMFAPMGEDRDMGAILDDLCRQMADALPEIETP* |
| AlbB | MHHHHHHHHGGGSGGG*SENLYFQG*NPGETVLPPQLREEIALLAVYLLSSGRGLLEEPADYGIYRCTDGARRALQLLDEHGGSTARLTAVRERLDEVMFAPMGEDRDMGAILDDLCRQMADALPEIETP* |
| AlbC | MHHHHHHHHGGGSGGG*SENLYFQG*LAGLVPAPDHGMREEILGDRSRLIRQRGEHALIGISAGNSYFSQKNTVMLLQWAGQRFERTDVVYVDTHIDEMLIADGRSAQEAERSVKRTLKDLRRRLRRSLESVGDHAERFRVRSLSELQETPEYRAVRERTDRAFEEDAEFATACEDMVRAVVMNRPGDGVGISAEHLRAGLNYVLAEAPLFADSPGVFSVPSSVLCYHIDTPITAFLSRRETGFRAAEGQAYVVVRPQELADAA* |

[1]Stops are represented by asterisks.

[2]Underlined amino acids represent expression tags.

[3]Italicized amino acids represent TEV protease site.
